# Serotonin Reuptake Inhibition Reduces NMDAR-Associated Reward Liability Without Compromising Rapid Antidepressant Efficacy

**DOI:** 10.64898/2026.11.01.763059

**Authors:** Libiao Pan, Guan Wang, Jing Ji, Yuting Cai, Yingting Pang, Yanmei Chen, Qiaohui Lu, Xuhua Tang, Zhengtao Hu, Baiyang Zheng, Yangyang Xiang, Qiongqiong Hou, Ke Nie, Jiangyu Zhao, Tianwen Hu, Yang He, Yan Zhang, Vivien Shen, Weiliang Zhu, Haji Akber Aisa, Guanghui Tian, Jingshan Shen

## Abstract

Ketamine has changed the treatment landscape for depression by providing rapid antidepressant effects, yet its clinical utility remains limited by psychotomimetic effects and abuse potential. It remains unclear whether rapid antidepressant efficacy can be dissociated from mesolimbic dopamine activation and its associated reward-related effects. Here, we show that the serotonin reuptake inhibitor escitalopram, by enhancing serotonergic signaling, attenuates S-ketamine-induced dopamine (DA) release in the nucleus accumbens (NAc). Inspired by this finding, we designed and evaluated VV147 by incorporating SERT inhibition into NMDAR antagonism to reduce the reward-related liability of NMDAR blockade. Unlike S-ketamine, VV147 did not increase NAc dopamine signaling but retained rapid antidepressant effects in chronic stress models of depression. Together, these findings suggest a potential dissociation between rapid antidepressant efficacy and acute dopamine activation, providing a possible strategy for developing rapid-acting antidepressants with reduced dopamine-related reward effects.

## Introduction

Major depressive disorder (MDD) is a leading cause of disability worldwide, and the delayed onset of action of conventional antidepressants remains a major challenge, particularly for patients with severe or treatment-resistant depression^1–3^. The discovery that ketamine can produce rapid and sustained antidepressant effects has transformed the search for faster-acting treatments and established NMDARs as an important target for antidepressant development^1,4–6^. However, the clinical use of ketamine is limited by acute dissociative and psychotomimetic effects, as well as concerns about its abuse potential^5,7,8^. These limitations raise a question: can rapid antidepressant efficacy be retained while reducing ketamine-associated neurobehavioral adverse effects? A better understanding of the mechanisms underlying ketamine’s therapeutic and undesirable effects is needed to address this question. Dissociative and psychotomimetic effects have been associated with NMDAR antagonism, although their relationship to the extent and pattern of receptor blockade remains incompletely understood^9,10^. It is also unclear whether the neural processes underlying ketamine’s reward-related effects can be modulated independently of its antidepressant actions. The mesolimbic dopamine system, particularly DA signaling in the NAc, plays a central role in reward processing and reinforcement^11–13^. Ketamine can alter mesolimbic DA transmission, potentially contributing to its reward-related effects and abuse liability^4,13^. Understanding how these DA responses are regulated may help guide the development of rapid-acting antidepressants with a more favorable neurobehavioral profile. Previous studies have shown that, beyond its established role in mood regulation, serotonergic signaling modulates mesolimbic DA transmission through multiple receptor subtypes^14,15,16^. These findings suggest that enhancing serotonergic signaling may help limit excessive drug-induced DA responses. We therefore examined whether escitalopram, a selective serotonin reuptake inhibitor (SSRI) that enhances serotonergic signaling by blocking 5-HT reuptake, could attenuate S-ketamine-induced DA release in NAc, and further investigated the receptor mechanisms involved. We next asked whether serotonin reuptake inhibition could reduce NMDAR blockade-associated reward-related effects while preserving rapid antidepressant activity. Here, *in vivo* recordings showed that escitalopram attenuated S-ketamine-induced dopamine release in the NAc. Inspired with this finding, we hypothesized that incorporating SERT inhibition into NMDAR antagonism could preserve rapid antidepressant efficacy while reducing dopamine-related reward effects. Rational drug design and structural optimization were starting from 3-methoxymorphinan (3-MM), which was chosen as a lead scaffold due to its initial bioactivities towards both targets of SERT and NMDAR. The rational drug design practiced the idea of “small change, big benefit” and yielded VV147, an NMDAR antagonist with SERT inhibitory activity. Unlike S-ketamine, VV147 did not increase NAc DA signaling, induce conditioned place preference (CPP), or produce locomotor hyperactivity or impaired rotarod performance. Nevertheless, VV147 produced rapid antidepressant effects in both chronic unpredictable mild stress (CUMS) and chronic restraint stress models (CRS). Together, these findings suggest that introducing SERT inhibition into NMDAR antagonism may provide a strategy for developing rapid-acting antidepressants with reduced dopamine-related reward effects.

## Results

### Escitalopram attenuates S-ketamine-induced dopamine responses in the NAc

We first examined the effects of S-ketamine on dopamine signaling in the NAc using fiber photometry. C57BL/6J mice received stereotaxic injections of an AAV encoding the genetically encoded dopamine sensor GRAB-DA into the NAc and were allowed to recover for 3 weeks before testing^17^. S-ketamine (30 mg/kg, i.p.) produced a robust increase in the GRAB-DA fluorescence signal. Cocaine (15 mg/kg, i.p.), used as a positive control, also markedly increased the signal (Fig. 1a–c). In the CPP test, S-ketamine induced a significant preference for the drug-paired chamber (Fig. 1d).

**Figure 1.**
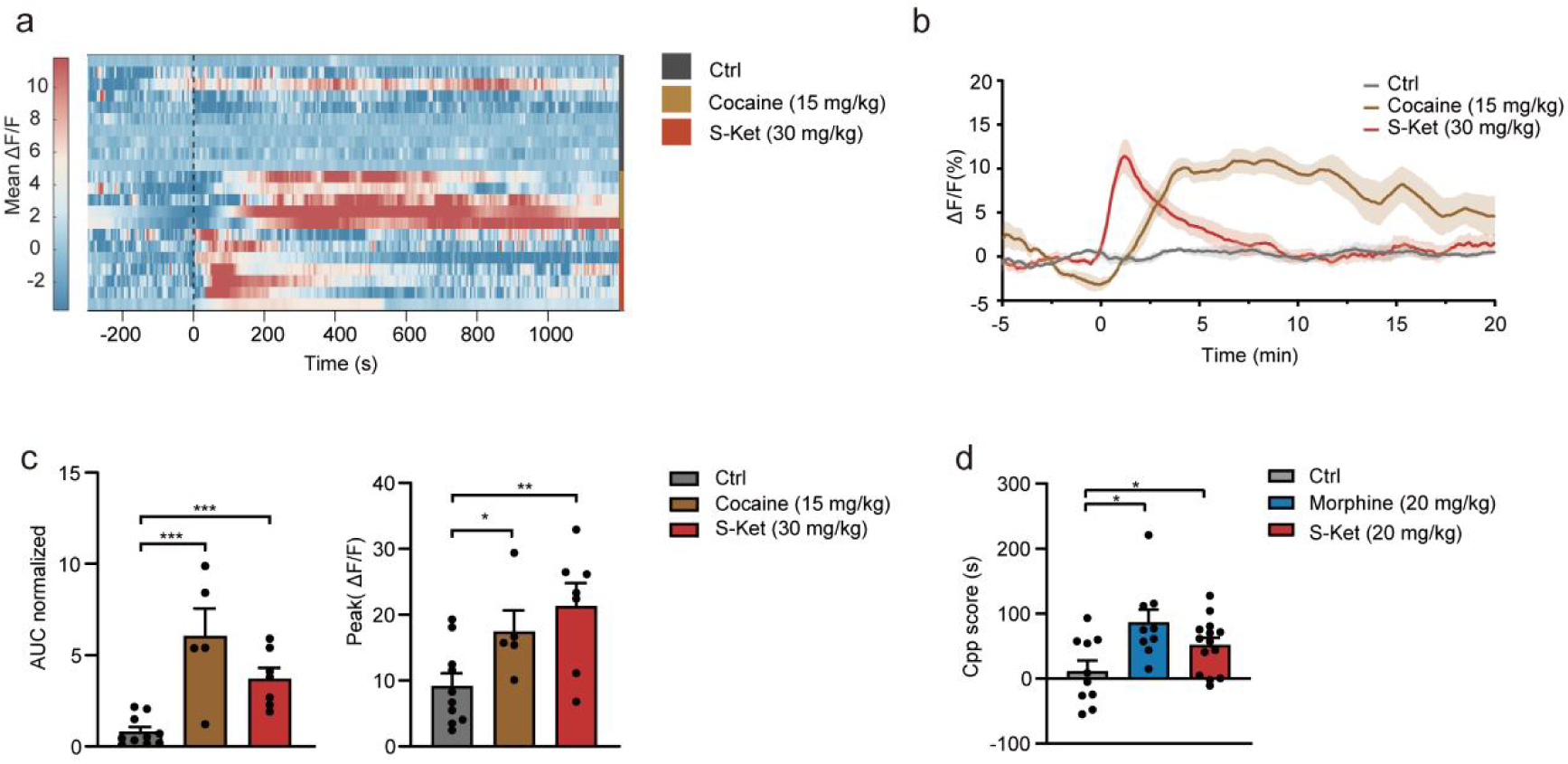
S-ketamine increases NAc dopamine signaling and induces conditioned place preference. (**a, b**) Normalized NAc DA dynamics in response to i.p. injections of saline (Ctrl), Cocaine (15 mg/kg), and S-ketamine (S-Ket) (30 mg/kg), depicted as single-trial heatmaps (a) and averaged traces (b). (**c**) Quantification of signal magnitude. Left, normalized area under the curve (AUC). Right, peak fluorescence change (Peak ΔF/F0). For AUC, Ctrl vs. Cocaine (15 mg/kg), unpaired t test, P =0.0033; Ctrl vs. S-Ket (30 mg/kg), unpaired t test, P =0.0001. For Peak fluorescence change, Ctrl vs. Cocaine (15 mg/kg), unpaired t test, P =0.0340; Ctrl vs. S-Ket (30 mg/kg), unpaired t test, P =0.0047. Ctrl, n=10; Cocaine (15 mg/kg), n=5; S-Ket (30 mg/kg), n=7. (**d**) Conditioned place preference induced by morphine and S-ketamine. Ctrl vs. Morphine (20 mg/kg), Mann-Whitney test, P = 0.0101; Ctrl vs. S-Ket (20 mg/kg), unpaired t test, P = 0.0460. Ctrl, n = 10; Morphine (20 mg/kg), n = 10; S-Ket (20 mg/kg), n = 15. *P < 0.05; **P < 0.01; ***P < 0.001. Error bar, SEM.

We next examined whether escitalopram could modify the dopamine response to S-ketamine. Escitalopram (10, 30, or 60 mg/kg, i.p.) was administered before S-ketamine. The 10 mg/kg dose had no significant effect on the S-ketamine-induced fluorescence response. At 30 mg/kg, escitalopram produced a downward trend, whereas 60 mg/kg significantly attenuated the response (Fig. 2a–c).

**Figure 2.**
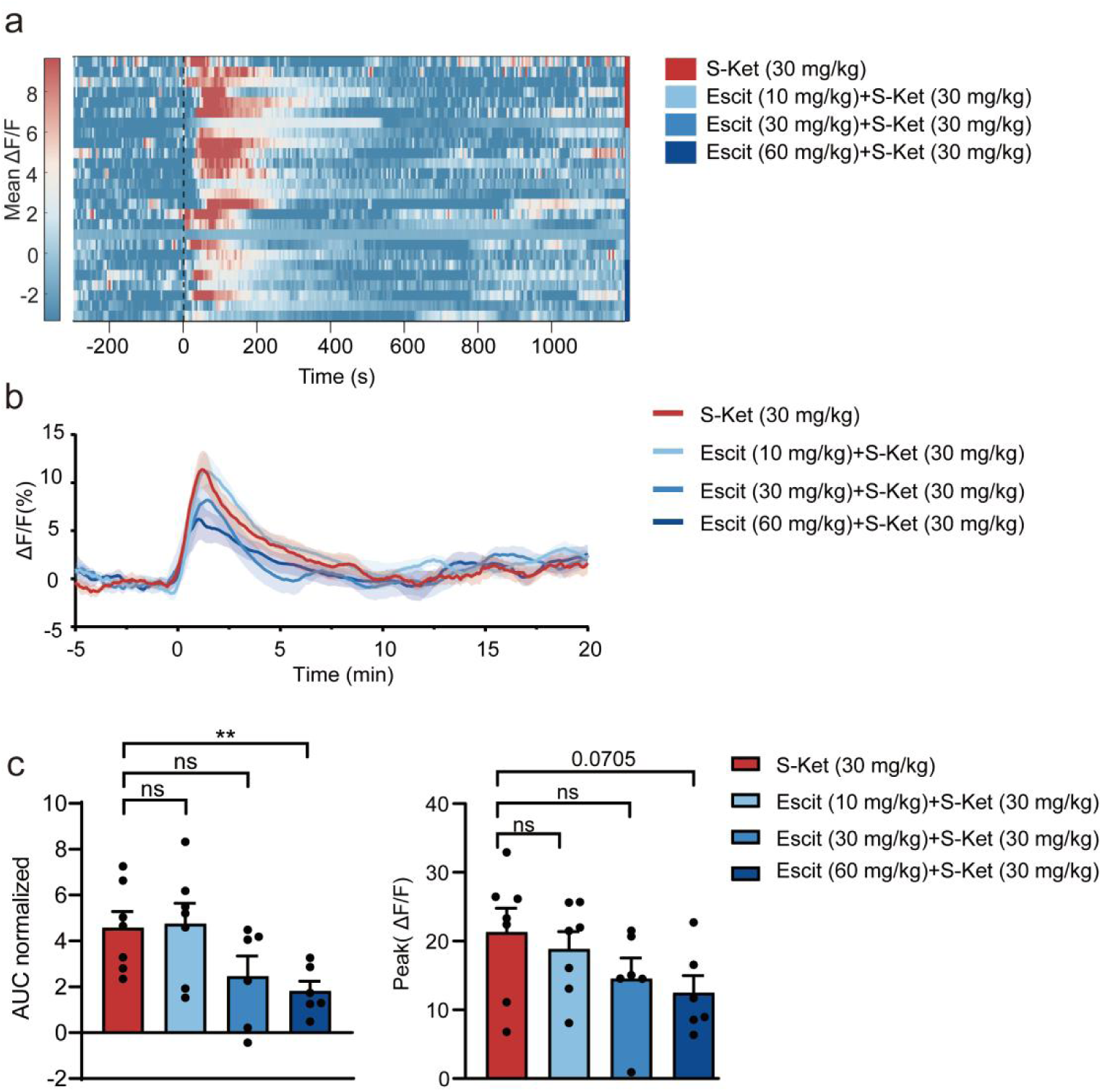
Escitalopram attenuates S-ketamine-induced NAc dopamine signaling. (a, **b)** Normalized NAc DA dynamics in response to i.p. injections of S-Ket (30 mg/kg) and varying doses of Escitalopram (Escit) (10, 30, or 60 mg/kg), depicted as single-trial heatmap (a) and averaged trace (b). (c) Quantification of signal magnitude. Left: Normalized area under the curve (AUC). Right: Peak fluorescence change (Peak ΔF/F0). For AUC, Escit (10 mg/kg) + S-Ket (30 mg/kg) vs. S-Ket (30 mg/kg), unpaired t test, P = 0.8794; Escit (30 mg/kg) + S-Ket (30 mg/kg) vs. S-Ket (30 mg/kg), unpaired t test, P = 0.0860; Escit (60 mg/kg) + S-Ket (30 mg/kg) vs. S-Ket (30 mg/kg), unpaired t test, P = 0.0090. For Peak fluorescence change, Escit (10 mg/kg) + S-Ket (30 mg/kg) vs. S-Ket (30 mg/kg), unpaired t test, P = 0.5788; Escit (30 mg/kg) + S-Ket (30 mg/kg) vs. S-Ket (30 mg/kg), unpaired t test, P = 0.1760; Escit (60 mg/kg) + S-Ket (30 mg/kg) vs. S-Ket (30 mg/kg), unpaired t test, P = 0.0705. S-Ket (30 mg/kg), n = 7, Escit (10 mg/kg) + S-Ket (30 mg/kg), n = 7, Escit (30 mg/kg) + S-Ket (30 mg/kg), n = 6, Escit (60 mg/kg) + S-Ket (30 mg/kg), n = 6. **P < 0.01; ns, not significant. Error bar, SEM.

These data indicate that serotonin reuptake inhibition can attenuate S-ketamine-induced dopamine signaling in the NAc.

### 5-HT1B and 5-HT_2C_ receptors contribute to serotonergic suppression of S-ketamine-induced dopamine release

We next investigated which serotonin receptor subtypes contribute to the modulation of S-ketamine-induced dopamine signaling. Given the established roles of 5-HT_1B_ and 5-HT_2C_ receptors in regulating mesolimbic dopamine neurotransmission^18^, we examined whether activation of either receptor could attenuate the dopamine response to S-ketamine. Administration of the 5-HT_1B_ receptor agonist CP-93129 (2 mg/kg, i.p.) or the 5-HT_2C_ receptor agonist Ro60-0175 (3 mg/kg, i.p.) significantly attenuated the S-ketamine-induced increase in the NAc GRAB-DA fluorescence signal (Fig. 3a– c).

**Figure 3.**
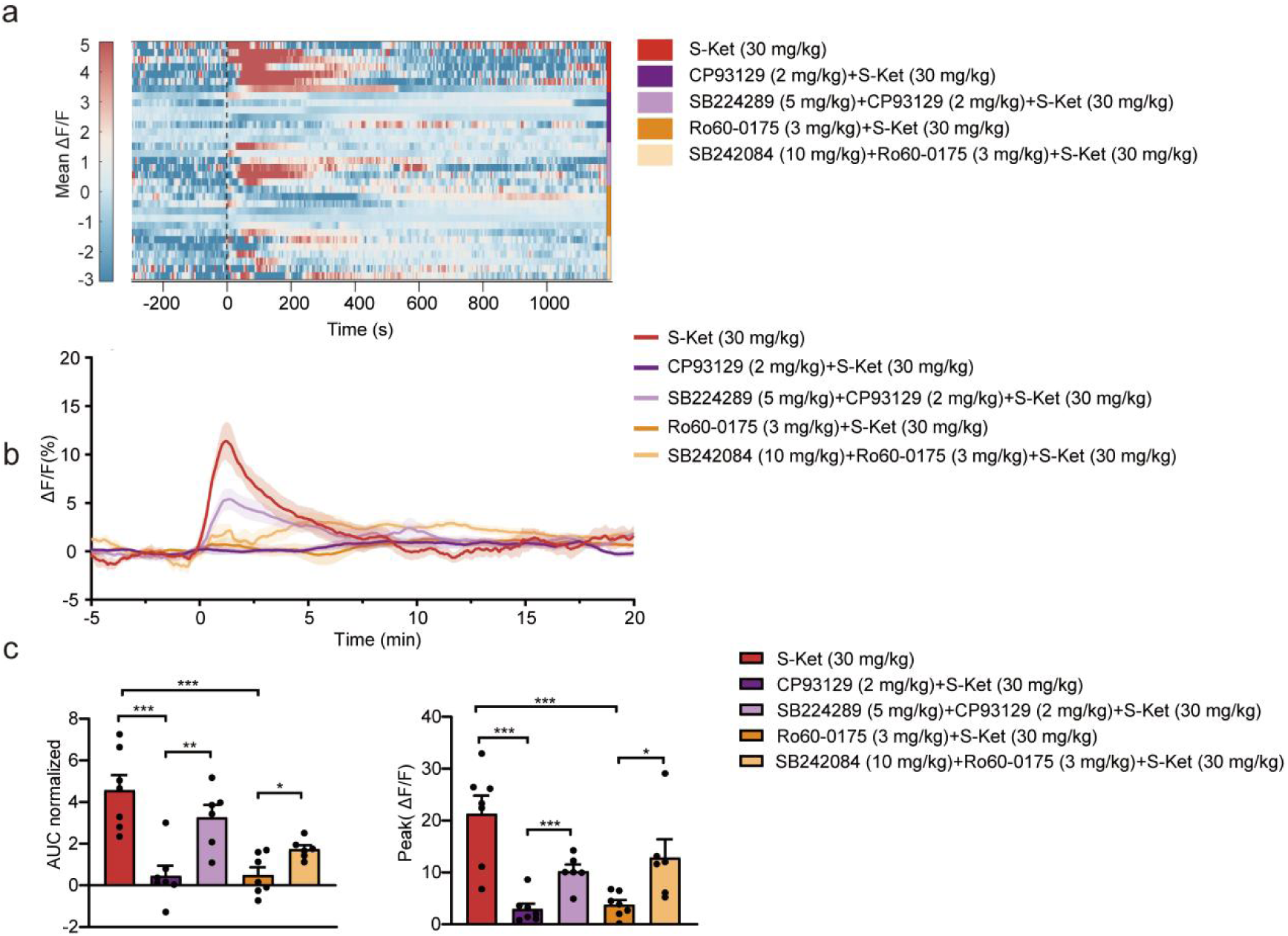
5-HT1B and 5-HT2C receptors regulate S-ketamine-induced NAc dopamine signaling. (a, **b)** 5-HT1B and 5-HT2C agonist/antagonist delivery evokes NAc gDA dynamics (normalized), depicted as heatmap (a) and mean trace (b). (c) AUC (left) and peak (right) of DA signals. For AUC, CP93129 (2 mg/kg) + S-Ket (30 mg/kg) vs. S-Ket (30 mg/kg), unpaired t test, P = 0.0005; Ro60-0175 (3 mg/kg) + S-Ket (30 mg/kg) vs. S-Ket (30 mg/kg), unpaired t test, P = 0.0003; SB224289 (5 mg/kg) + CP93129 (2 mg/kg) + S-Ket (30 mg/kg) vs. CP93129 (2 mg/kg) + S-Ket (30 mg/kg), unpaired t test, P = 0.0036; SB242084 (10 mg/kg) + Ro60-0175 (3 mg/kg) + S-Ket (30 mg/kg) vs. Ro60-0175 (3 mg/kg) + S-Ket (30 mg/kg), unpaired t test, P = 0.0156. For peak, CP93129 (2 mg/kg) + S-Ket (30 mg/kg) vs. S-Ket (30 mg/kg), unpaired t test, P = 0.0003; Ro60-0175 (3 mg/kg) + S-Ket (30 mg/kg) vs. S-Ket (30 mg/kg), unpaired t test, P = 0.0004; SB224289 (5 mg/kg) + CP93129 (2 mg/kg) + S-Ket (30 mg/kg) vs. CP93129 (2 mg/kg) + S-Ket (30 mg/kg), unpaired t test, P = 0.0009; SB242084 (10 mg/kg) + Ro60-0175 (3 mg/kg) + S-Ket (30 mg/kg) vs. Ro60-0175 (3 mg/kg) + S-Ket (30 mg/kg), unpaired t test, P = 0.0209. S-Ket (30 mg/kg), n = 7; CP93129 (2 mg/kg) + S-Ket (30 mg/kg), n = 7; Ro60-0175 (3 mg/kg) + S-Ket (30 mg/kg), n = 7; SB224289 (5 mg/kg) + CP93129 (2 mg/kg) + S-Ket (30 mg/kg), n = 6; SB242084 (10 mg/kg) + Ro60-0175 (3 mg/kg) + S-Ket (30 mg/kg), n = 6. *P < 0.05; **P < 0.01; ***P < 0.001. Error bar, SEM.

To further assess the involvement of these receptors, we pretreated mice with the corresponding selective antagonists before administration of the agonists and S-ketamine. Pretreatment with the 5-HT_1B_ receptor antagonist SB224289 (5 mg/kg, i.p.) or the 5-HT_2C_ receptor antagonist SB242084 (10 mg/kg, i.p.) partially reversed the agonist-induced attenuation of the S-ketamine-evoked NAc GRAB-DA fluorescence response (Fig. 3a–c). These findings support the involvement of both 5-HT_1B_ and 5-HT_2C_ receptors in the serotonergic modulation of S-ketamine-induced dopamine signaling in the NAc.

### Drug design and evaluation of VV147: small change, big benefit

Inspired with the finding that serotonin reuptake inhibition attenuated the dopamine response induced by S-ketamine, we sought to incorporate SERT inhibition into an NMDAR antagonist to possess rapid antidepressant efficacy while reducing NMDAR antagonist-associated reward liability.

3-Methoxymorphinan (3-MM), the N-demethylated metabolite of the antitussive agent dextromethorphan, was chosen from our compound library as a lead scaffold for its initial inhibitory activity towards both NMDAR and SERT, aiming to enhance its moderate SERT inhibitory activity while maintaining an appropriate active level of NMDAR inhibition and further, improving metabolic stability. Molecular docking of 3-MM with SERT protein suggested that the 2-position is sterically accessible, permitting further derivatization for testing of SERT activity, and introducing substituent on this position may also impose steric hindrance therefor to impede enzyme binding to the 3-methylhydroxy group for improving its metabolic stability of the designed compound, and that is a well-established rational drug design strategy in medicinal chemistry. The substituents introduced on to this 2-position were chloro, bromo, and alkyl groups, affording compounds **2 – 9** respectively. *In vitro* assays demonstrated that the 2-substituented compounds of chloro, bromo, methyl, and ethyl (compound **2**, **3**, **4** and **5**) enhanced SERT inhibition to varying degrees. Among these compounds, compound **4** exhibited the most potent SERT inhibition, with an IC₅₀ value of 190 nM (Table 1). Although the introducing of 2-substituents moderately attenuated NMDAR activity relative to 3-MM, the compounds retained micromolar IC₅₀ values. The metabolic stability of compounds **2 – 5** was then evaluated using human liver microsomes (HLMs). Remarkably, after 60 min of incubation, Compound **4** displayed the highest stability, substantially outperforming 3-MM under identical conditions. Moreover, the percentage of parent compound remaining for Compound **4** did not decrease significantly over the incubation period, precluding the calculation of both half-life and clearance. This low-clearance profile strongly suggests that the compound is metabolized extremely slowly in vitro, demonstrating relatively good metabolic stability. To further increase the metabolic stability of Compound **4**, we employed a deuteration strategy to generate Compound **10** (VV147, Table 1). VV147 retained target potency comparable to that of Compound **4** while exhibiting increased metabolic stability in human liver microsomes.

**Table 1.**
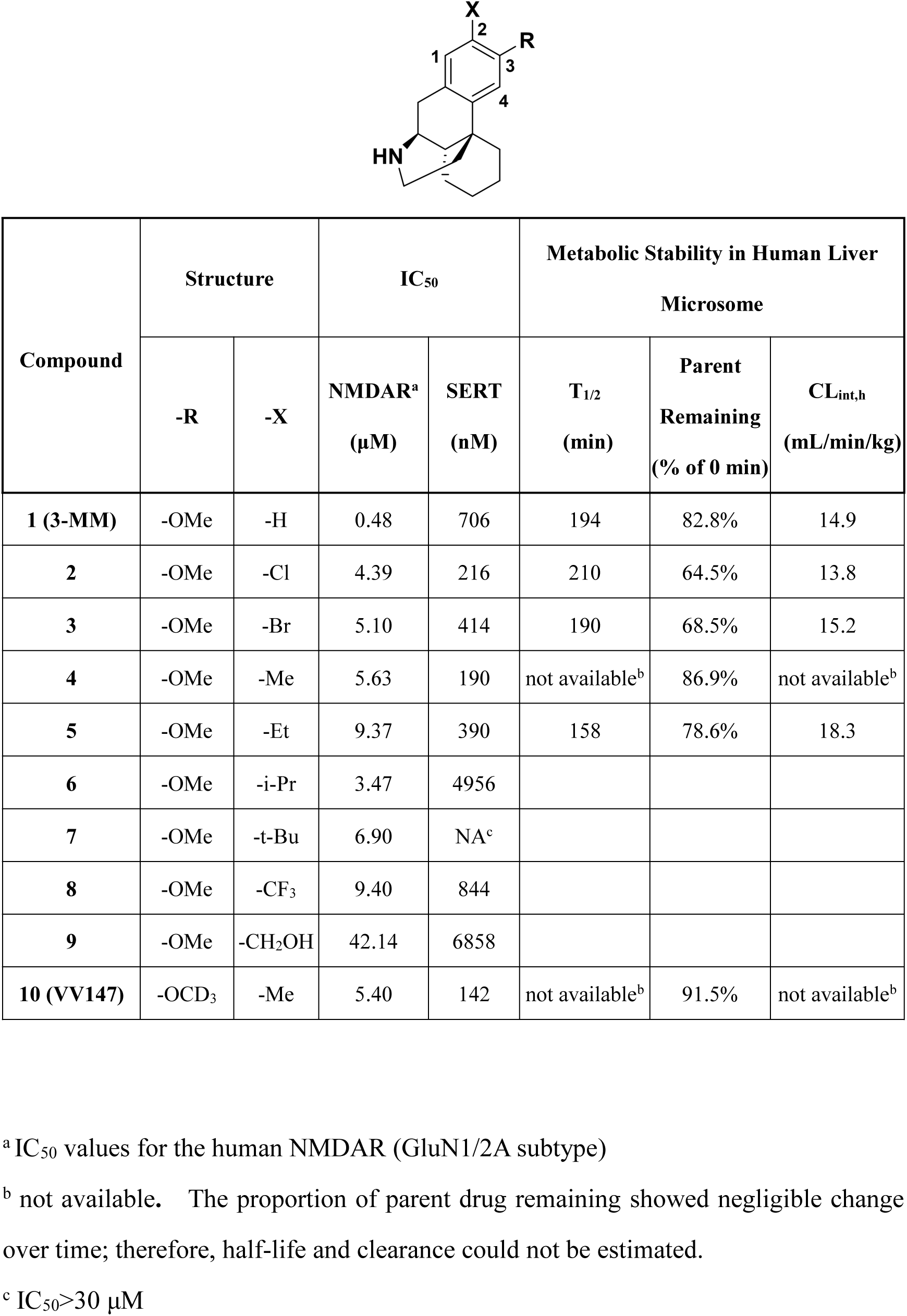
Structure and activities of compounds 1–10 against NMDAR, SERT and Metabolic Stability in Human Liver Microsome.

| **Compound** | **Structure** |  | **IC<sub>50</sub>** |  | **Metabolic Stability in Human Liver Microsome** |  |  |
|---|---|---|---|---|---|---|---|
|  | **-R** | **-X** | **NMDAR<sup>a</sup><br>(µM)** | **SERT<br>(nM)** | **T<sub>1/2</sub><br>(min)** | **Parent Remaining<br>(% of 0 min)** | **CL<sub>int,h</sub><br>(mL/min/kg)** |
| **1 (3-MM)** | -OMe | -H | 0.48 | 706 | 194 | 82.8% | 14.9 |
| **2** | -OMe | -Cl | 4.39 | 216 | 210 | 64.5% | 13.8 |
| **3** | -OMe | -Br | 5.10 | 414 | 190 | 68.5% | 15.2 |
| **4** | -OMe | -Me | 5.63 | 190 | not available<sup>b</sup> | 86.9% | not available<sup>b</sup> |
| **5** | -OMe | -Et | 9.37 | 390 | 158 | 78.6% | 18.3 |
| **6** | -OMe | -i-Pr | 3.47 | 4956 |  |  |  |
| **7** | -OMe | -t-Bu | 6.90 | NA<sup>c</sup> |  |  |  |
| **8** | -OMe | -CF<sub>3</sub> | 9.40 | 844 |  |  |  |
| **9** | -OMe | -CH<sub>2</sub>OH | 42.14 | 6858 |  |  |  |
| **10 (VV147)** | -OCD<sub>3</sub> | -Me | 5.40 | 142 | not available<sup>b</sup> | 91.5% | not available<sup>b</sup> |

<sup>a</sup> IC<sub>50</sub> values for the human NMDAR (GluN1/2A subtype)
<sup>b</sup> not available**.** The proportion of parent drug remaining showed negligible change over time; therefore, half-life and clearance could not be estimated.
<sup>c</sup> IC<sub>50</sub>>30 µM

In summary, this process demonstrates that small change of a chemical structure with rational drug design strategy can make a big difference in drug-like property optimization regarding to target potency and metabolic stability, ultimately yielding VV147 as a promising candidate for further druggability evaluation.

### VV147 does not elevate NAc dopamine or induce conditioned reward

VV147 inhibited both SERT and NMDARs, consistent with its dual-target profile (Fig. 4a,b). We next examined its effects on mesolimbic dopamine signaling and reward-related behavior. VV147 did not produce a detectable increase in NAc GRAB-DA fluorescence signal (Fig. 4c–e). Similarly, unlike S-ketamine, VV147 did not induce conditioned place preference (Fig. 4f).

**Figure 4.**
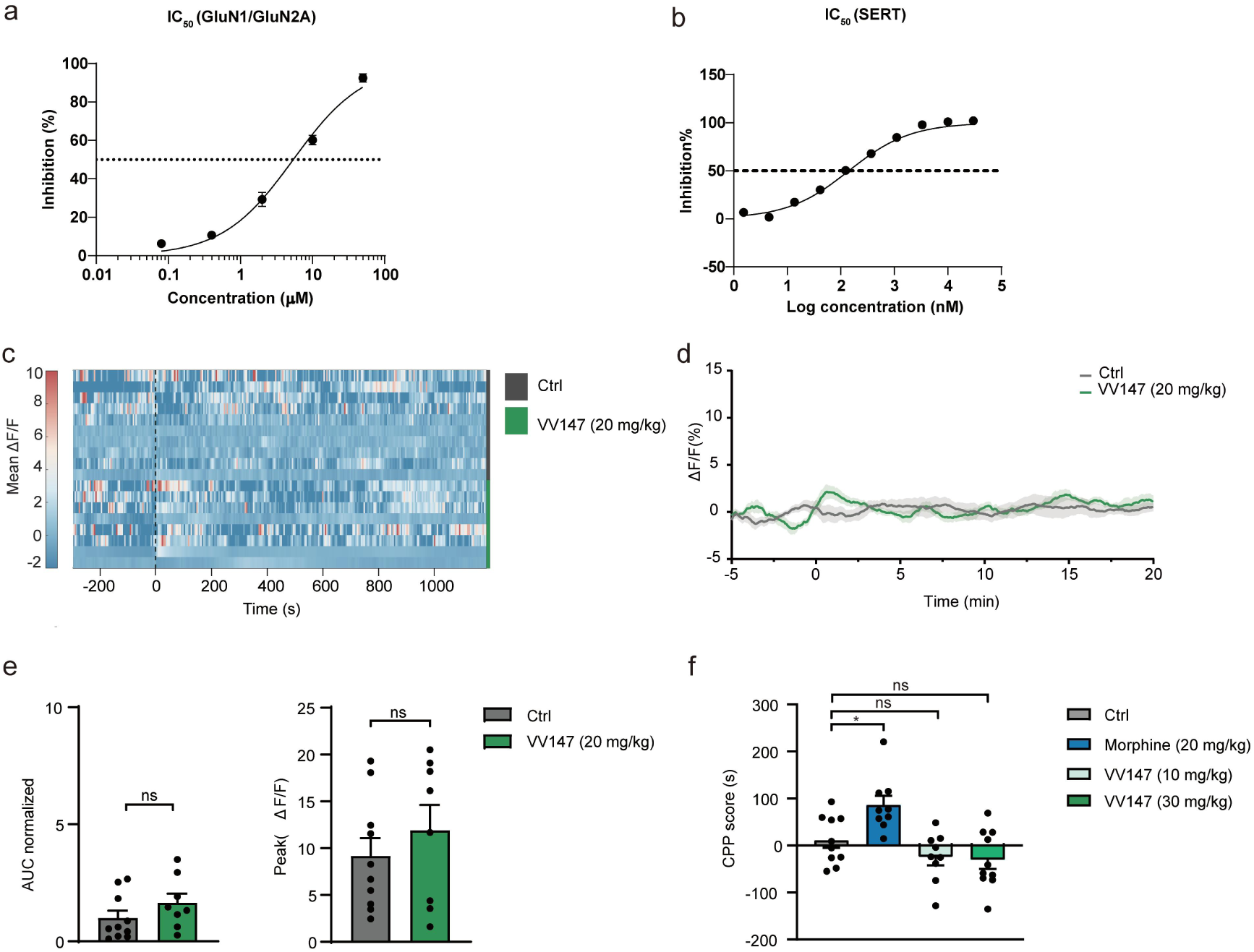
VV147 acts as a dual NMDAR and SERT modulator without increasing NAc dopamine signaling or inducing conditioned place preference. **(a)** Concentration-response curves of VV147 in HEK293 cells transfected with GluN1/GluN2A receptors. (b) Concentration-response curves of VV147 in HEK293 cells stably expressing human SERT. (c, d) VV147 (20 mg/kg) administration-evoked response dynamics: heatmap (c) and mean trace (d). (e) Response amplitude: normalized AUC (left) and peak (right). For AUC, VV147 (20 mg/kg) vs. Ctrl, unpaired t test, P = 0.2065. For peak, VV147 (20 mg/kg) vs. Ctrl, unpaired t test, P = 0.4112. Ctrl, n = 10; VV147 (20 mg/kg), n = 8. (f) Conditioned place preference induced by morphine but not VV147. Ctrl vs. Morphine (20 mg/kg), Mann-Whitney test, P = 0.0101; Ctrl vs. VV147 (10 mg/kg), unpaired t test, P = 0.1509. Ctrl vs. VV147 (30 mg/kg), unpaired t test, P = 0.1205. n = 10 per group. *P < 0.05; ns, not significant. Error bar, SEM.

We then investigated whether VV147 could attenuate the dopamine response to S-ketamine. VV147 (20 or 40 mg/kg, i.p.) was administered 30 min before S-ketamine. Pretreatment with VV147 at 40 mg/kg reduced the S-ketamine-induced increase in NAc GRAB-DA fluorescence, whereas the 20 mg/kg dose did not produce a comparable effect (Fig. 5a–c).

**Figure 5.**
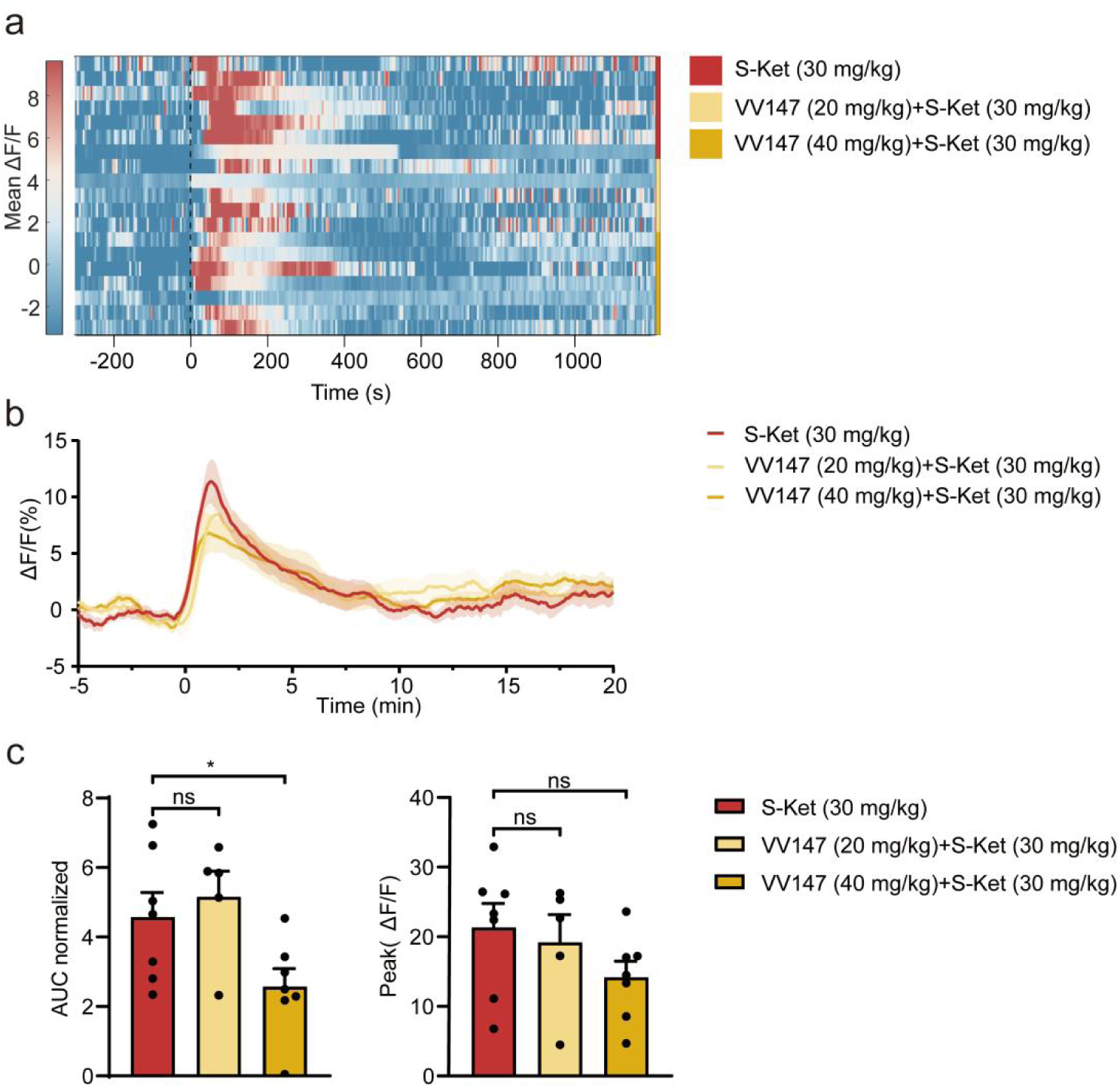
VV147 attenuates S-ketamine-induced NAc dopamine release. (a, **b)** Normalized NAc DA dynamics evoked by i.p. S-Ket (30 mg/kg) and varying VV147 doses (20 or 40 mg/kg): heatmap (a) and averaged trace (b). (c) Quantification of response AUC (left) and peak (right) of DA signals to different dose groups. For AUC, VV147 (20 mg/kg) + S-Ket (30 mg/kg) vs. S-Ket (30 mg/kg), unpaired t test, P = 0.5927; VV147 (40 mg/kg) + S-Ket (30 mg/kg) vs. S-Ket (30 mg/kg), unpaired t test, P = 0.0424. For peak, VV147 (20 mg/kg) + S-Ket (30 mg/kg) vs. S-Ket (30 mg/kg), unpaired t test, P = 0.7004; VV147 (40 mg/kg) + S-Ket (30 mg/kg) vs. S-Ket (30 mg/kg), unpaired t test, P = 0.112. S-Ket (30 mg/kg), n = 7; VV147 (20 mg/kg) + S-Ket (30 mg/kg), n = 5; VV147 (40 mg/kg) + S-Ket (30 mg/kg), n = 7. *P < 0.05; ns, not significant. Error bar, SEM.

Together, these results demonstrate that VV147 inhibits both SERT and NMDARs without increasing NAc dopamine signaling or inducing CPP.

### VV147 produces rapid antidepressant effects that do not require potent SERT inhibition

Having characterized the dopaminergic and reward-related profile of VV147, we next evaluated its antidepressant efficacy in two chronic stress models: chronic unpredictable mild stress (CUMS) and chronic restraint stress (CRS). In both models, a single administration of VV147 significantly increased sucrose preference and reduced immobility time in the forced swim test. These effects were similar in direction to those observed following S-ketamine treatment, supporting the rapid antidepressant effects of VV147 in chronic stress models (Fig. 6a-c).

**Figure 6.**
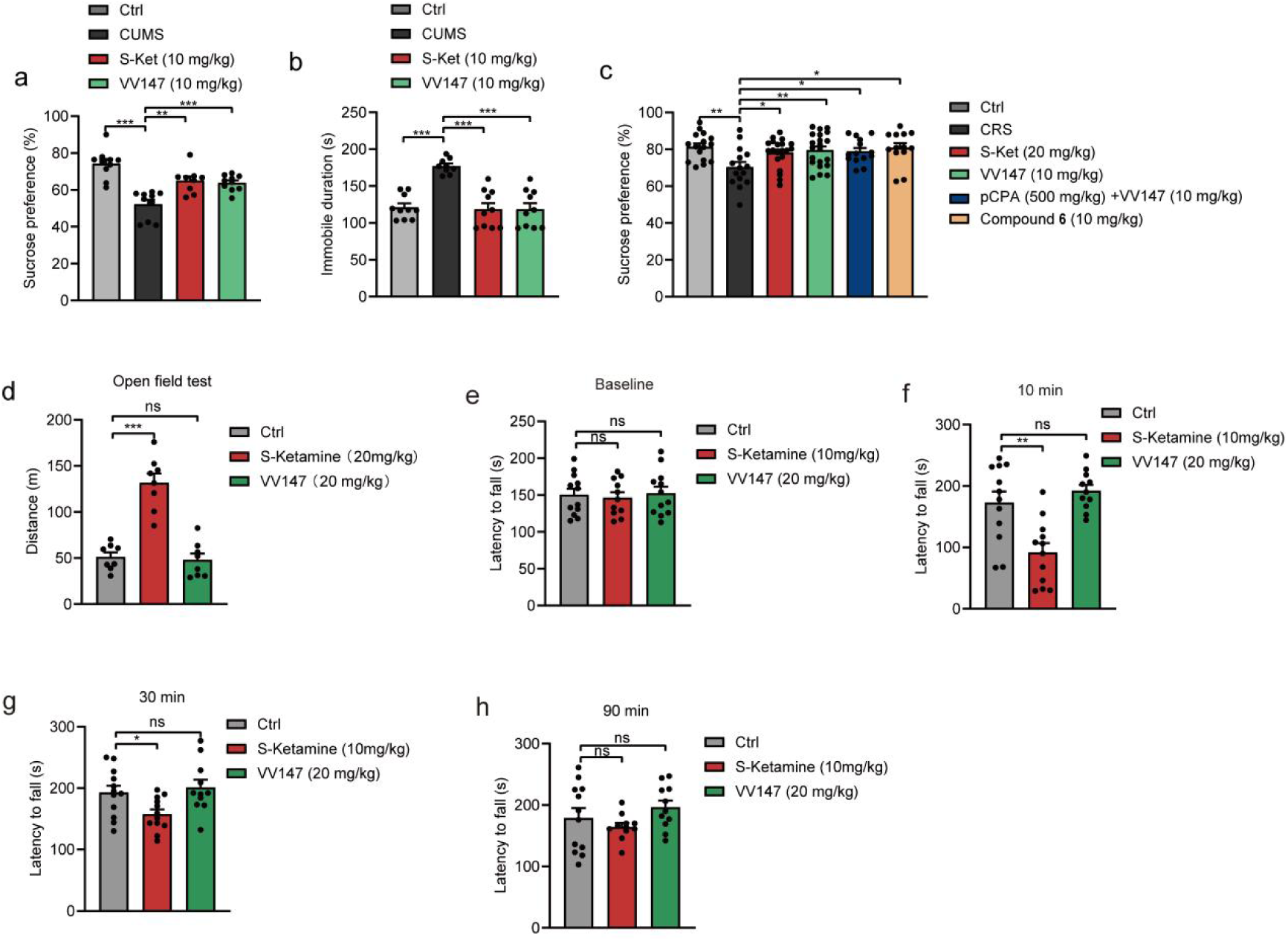
VV147 produces rapid antidepressant effects independent of potent SERT inhibition. (a-b) Sucrose preference and duration of immobility was assessed in CUMS-exposed mice with SPT and FST respectively. (a) Sucrose preference. Ctrl vs. CUMS, Mann-Whitney test, P < 0.0001; CUMS vs. S-Ket (10 mg/kg), Mann-Whitney test, P = 0.0020; CUMS vs. VV147 (10 mg/kg), Mann-Whitney test, P = 0.0002. Ctrl, n = 10; CUMS, n=10; S-Ket (10 mg/kg), n = 9; VV147 (10 mg/kg), n = 10. (b) Immobile duration. Ctrl vs. CUMS, unpaired t test, P < 0.0001; CUMS vs. S-Ket (20 mg/kg), Welch’s t test, P < 0.0001; CUMS vs. VV147 (10 mg/kg), Welch’s t test, P < 0.0001. Ctrl, n = 10; CUMS, n=9; S-Ket (20 mg/kg), n = 10; VV147 (10 mg/kg), n = 10. (c) Sucrose preference was assessed in CRS-exposed mice with SPT. Ctrl vs. CRS, unpaired t test, P = 0.0022; CRS vs. S-Ket (20 mg/kg), Mann-Whitney test, P = 0.0494; CRS vs. VV147 (10 mg/kg), unpaired t test, P = 0.0081; CRS vs. pCPA (500 mg/kg) +VV147 (10 mg/kg), unpaired t test, P = 0.0164; CRS vs. Compound 6 (10 mg/kg), unpaired t test, P = 0.0100. Ctrl, n = 17; CRS, n=15; S-Ket (20 mg/kg), n = 20; VV147 (10 mg/kg), n = 21; pCPA (500 mg/kg) + VV147 (10 mg/kg), n = 14; Compound 6 (10 mg/kg), n = 13. (d) Distance travelled after i.p. injections in open field test. Ctrl vs. S-Ket, unpaired t test, P < 0.0001; Ctrl vs. VV147, unpaired t test, P = 0.6952. n = 8 per group. (e-h) Latency to fall before and after i.p. injections in rotarod test. (e) Baseline. Ctrl vs. S-Ket, unpaired t test, P = 0.7224; Ctrl vs. VV147, unpaired t test, P = 0.8581. Ctrl, n = 12; S-Ket, n = 11; VV147, n = 12. (f) Latency (10 min post-injection). Ctrl vs. S-Ket, unpaired t test, P = 0.0025; Ctrl vs. VV147, Welch’s t test, P = 0.3585. Ctrl, n = 12; S-Ket, n = 12; VV147, n = 11. (g) Latency (30 min post-injection). Ctrl vs. S-Ket, unpaired t test, P = 0.0175; Ctrl vs. VV147, unpaired t test, P = 0.6245. Ctrl, n = 12; S-Ket, n = 12; VV147, n = 11.(h) Latency (90 min post-injection). Ctrl vs. S-Ket, Welch’s t test, P = 0.4044; Ctrl vs. VV147, unpaired t test, P = 0.3911. Ctrl, n = 12; S-Ket, n = 11; VV147, n = 11. *P < 0.05; **P < 0.01; ***P < 0.001. Error bar, SEM.

We next investigated whether serotonergic signaling was required for the rapid antidepressant effects of VV147. We depleted brain serotonin using p-chlorophenylalanine (pCPA) and assessed behavioral responses in the CRS model^19^. Serotonin depletion did not significantly attenuate the effects of VV147 on sucrose preference in the CRS model (Fig. 6c). We also examined compound **6**, which retained NMDAR-inhibitory activity (IC₅₀ = 3.47 μM) but showed substantially weaker SERT-inhibitory activity (IC₅₀ = 4.96 μM) than VV147. Despite its weaker SERT-inhibitory activity, compound **6** also produced significant antidepressant effects in the behavioral assays (Fig. 6c).

Having established the rapid antidepressant efficacy of VV147, we next assessed its behavioral tolerability, focusing on adverse effects associated with ketamine^20–22^. In the open-field test, VV147 did not induce the locomotor hyperactivity observed following S-ketamine administration (Fig. 6d). Similarly, VV147 did not significantly affect rotarod performance, in contrast to the motor coordination deficits observed following S-ketamine administration (Fig. 6e-h). These differences may be related to the distinct NMDAR antagonistic properties of VV147 and S-ketamine, as ketamine-associated neuropsychiatric and motor adverse effects have been linked to NMDAR channel blockade^23^.

Together, these findings demonstrate the rapid antidepressant-like effects of VV147, suggest that potent SERT inhibition may not be necessary for these effects, and indicate that VV147 exhibits fewer motor side effects than S-ketamine.

## Discussion

This study examined whether escitalopram attenuated the S-ketamine-induced increase in NAc GRAB-DA fluorescence. This finding prompted the design of VV147, which adds SERT inhibition to reduce NMDAR-associated reward liability. Unlike S-ketamine, VV147 did not produce a detectable increase in NAc GRAB-DA fluorescence, induce CPP, or cause the locomotor hyperactivity and rotarod performance deficits observed after S-ketamine administration. Nevertheless, VV147 produced rapid antidepressant effects in chronic unpredictable mild stress and chronic restraint stress models. These effects were not significantly attenuated by serotonin depletion.

This suggests that SERT inhibition can attenuate ketamine-evoked dopamine responses in the NAc. Although dopamine transmission in this region is associated with reward-related and reinforcing effects, its relationship with antidepressant efficacy is less clear^14,17,18,24,25^. In the present study, VV147 produced rapid antidepressant-like effects without a detectable increase in NAc GRAB-DA fluorescence, suggesting that a detectable acute increase in NAc dopamine signaling is not necessary for the antidepressant effects of VV147. However, the absence of a detectable increase in fluorescence does not exclude more subtle changes in dopamine dynamics or responses occurring outside the measurement window. Taken together, the results indicate that the acute NAc dopamine response to an NMDAR antagonist alone may not reliably predict its antidepressant effects. These findings may inform the development of novel rapid-acting antidepressants targeting NMDARs alone or in combination with other molecular targets.

More broadly, our findings raise the possibility that SSRIs, such as escitalopram, fluoxetine, and paroxetine, as well as other drugs that enhance serotonergic signaling, may help mitigate the reward-related effects of drugs with abuse potential while preserving their desired pharmacological actions. Further preclinical and clinical studies are needed to determine whether and to what extent increasing serotonergic tone can reduce the abuse-related liability of rapid-acting antidepressants and other drugs with abuse potential.

Our pharmacological experiments further implicated 5-HT_1B_ and 5-HT_2C_ receptors in the regulation of S-ketamine-induced dopamine signaling. Activation of either receptor attenuated the dopamine response, while pharmacological blockade of 5-HT_1B_ and 5-HT_2C_ signaling partially reversed the attenuation. These findings support the involvement of both receptor systems, but do not establish that either receptor independently mediates the effect of escitalopram. The two receptors may regulate dopamine transmission through different mechanisms, including actions at dopamine terminals or within upstream neuronal circuits. The present experiments do not identify the relevant cellular sites of action or establish a direct pathway linking SERT inhibition to receptor activation. Further studies using receptor-selective interventions and cell-specific measurements will be needed to clarify how these receptor systems contribute to the serotonergic regulation of S-ketamine-evoked dopamine signaling and whether this regulation contributes to the therapeutic effects of ketamine.

Another notable finding was that serotonin depletion with p-CPA did not significantly attenuate the effect of VV147 on sucrose preference. Compound **6**, which has comparable NMDAR activity but substantially weaker SERT activity than VV147, also produced rapid antidepressant effects. These findings suggest that potent SERT inhibitory activity may not be necessary for the rapid antidepressant effects observed in the behavioral models examined. However, these results do not exclude a role for SERT in other aspects of VV147’s pharmacological profile, including its effects on monoaminergic signaling, the duration of its behavioral effects, or responses to repeated administration^26,27^. Whether SERT activity contributes to these effects requires further investigation.

Overall, VV147 produced rapid antidepressant effects but differed from S-ketamine in its neurochemical and behavioral profiles. Specifically, its antidepressant effects occurred without a detectable increase in NAc dopamine signaling. These findings underscore the need to determine the respective contributions of NMDAR and SERT modulation to the effects of VV147 and to link molecular-and circuit-level changes to behavioral outcomes. Determining the optimal balance between these two activities and defining their dose–response relationships will be important for optimizing the therapeutic index of this class of compounds and for balancing antidepressant efficacy against adverse effects. Whether the effects of VV147 observed in these preclinical models translate into clinical efficacy and predict its therapeutic potential in humans remains to be determined.

## MATERIALS AND METHODS

### Animals

Adult male C57BL/6J experimental mice were purchased from Vital River (Zhejiang, China) and Jihui (Shanghai, China). The mice were maintained on a 12-hour light/dark cycle (lights on at 07:00) at a constant temperature of 22–23°C and 40–60% humidity. Mice were randomly assigned to either the control or experimental groups. Before the experimental period, the animals were housed in the chambers within custom-designed stainless-steel cabinets. All experimental procedures were approved by the Ethics Committee of the Suzhou Institute of Materia Medica (Approval No.: 2024-11-lqh-01 and 2024-11-lqh-02).

### Stereotaxic surgery and viral delivery

For fiber photometry experiments, mice were deeply anesthetized with isoflurane (1.5% in pure oxygen) and positioned in a stereotaxic frame. Using a R-480 injection pump (RWD, China) at a flow rate of 30 nL/min, unilateral delivery of gDA (300 nL; pAAV-hSyn-DA3m-WPRE-hGH polyA^28^; titer 5.68E+12 vg/mL, PT-4720; BrainVTA(Wuhan) Co., Ltd, China) was targeted to the nucleus accumbens (NAc; AP +1.4 mm, ML ±1.2 mm, DV −4.0 mm relative to the dura). Following viral delivery, fiber-optic cannulae of appropriate length were implanted 0.1 mm dorsal to each injection site. Ferrule retention was achieved by sealing the exposed skull surface with tissue adhesive (1469SB; Vetbond, 3M, USA) and applying dental cement to the cannula side walls and surrounding skull. Prior to injection, a skull screw was placed 0.5 mm deep contralateral to the target site to enhance implant stability, and spacers were incorporated to minimize motion-induced artifacts. Fiber photometry recordings commenced 3 weeks after viral delivery.

For patch-clamp experiments, viruses encoding labels for glutamatergic neurons (200 nL; rAAV-CaMKIIα-mCherry-WPRE-hGH polyA; titer 5.28E+12 vg/mL, PT-0108; BrainVTA(Wuhan) Co., Ltd, China) and GABAergic neurons (200 nL; rAAV-GAD67-EGFP-WPRE; titer 5.57E+12 vg/mL, PT-12145; BrainVTA(Wuhan) Co., Ltd, China) were each injected into the mPFC (AP: +1.95 mm, ML: ± 0.33 mm, DV: −2.5 mm relative to the dura). Experiments were conducted 3 weeks after suturing.

### Fiber photometry recording and data analysis

*In vivo* fiber photometry was performed 3 weeks after viral injection and optic fiber implantation. A multi-channel fiber photometry system (RWD, China) was used to record dopamine dynamics in the NAc. The 470 nm and 410 nm LEDs were alternated at a camera frame rate of 60 fps to excite the dopamine sensor and the isosbestic control signal, respectively. The power at the tip of the optic fiber was adjusted to 20-30 µW.

Raw data were initially processed using the RWD software. The data preprocessing included background subtraction, and Exponential Fit for baseline correction. The processed data were then exported and further analyzed using custom-written scripts in MATLAB. The time of drug administration was defined as time 0. The mean fluorescence signal recorded during the 5-min period immediately before administration was defined as the baseline (*F_0_*). The dynamic changes in dopamine fluorescence were continuously recorded for 20-min post-administration. The Δ*F/F_0_* was calculated as (*F−F_0_*) / *F_0_* for the 20 min post-drug period to normalize the neural activity. For statistical analysis, the Δ*F/F_0_* values or the area under the curve (AUC) were quantified. Data were excluded if histological verification indicated viral expression or fiber placement outside the NAc.

### Whole-Cell Patch Clamping

HEK293T cells stably expressing human GluN2A were maintained in high-glucose Dulbecco’s modified Eagle’s medium (DMEM) supplemented with 10% fetal bovine serum and 1% penicillin–streptomycin at 37 °C in a humidified atmosphere containing 5% CO₂. Cells were seeded in 35-mm culture dishes and transiently transfected with an hGluN1-EGFP plasmid at 70–80% confluence using Lipofectamine 3000. For each dish, transfection complexes were prepared in Opti-MEM reduced-serum medium using 2.5 μg of plasmid DNA, 2.5 μL of Lipofectamine 3000, and 5 μL of P3000 reagent. The complexes were incubated for 15 min before addition to the cells. The culture medium was replaced 8 h after transfection, and cells were cultured for a further 24 h. Cells were subsequently dissociated, resuspended, counted, and seeded onto 10-mm coverslips in 24-well plates at 1 × 10⁵–3 × 10⁵ cells per well. Cells were allowed to adhere for 2 h before recording.

Test compounds were prepared as 100 mM stock solutions in dimethyl sulfoxide (DMSO) and diluted to the desired working concentrations in extracellular solution on the day of the experiment. The final DMSO concentration was maintained at 0.1% (v/v) in all compound-containing, vehicle control, and washout solutions. The extracellular solution contained (in mM): 140 NaCl, 2.5 KCl, 1 CaCl₂, 10 HEPES, and 20 sucrose, with the pH adjusted to 7.4 using NaOH. The pipette solution contained (in mM): 115 CsF, 10 CsCl, 10 EGTA, and 10 HEPES, with the pH adjusted to 7.2–7.3 using CsOH.

Whole-cell patch-clamp recordings were performed at room temperature (22–25°C) using an EPC-10 amplifier controlled by PatchMaster software (HEKA). Coverslips were transferred to a recording chamber and continuously perfused with extracellular solution. EGFP-positive cells were identified by fluorescence microscopy and selected for recording. After formation of a gigaohm seal and compensation of fast capacitance, whole-cell access was established. Cells were voltage-clamped at −70 mV, whole-cell capacitance was compensated, and series resistance compensation was set to 70%. Currents were acquired continuously in gap-free mode at a sampling rate of 20 kHz. After a stable baseline was established, GluN1/GluN2A receptor-mediated currents were evoked by co-application of glutamate and glycine, each at a final concentration of 100 μM. Once the evoked current reached a stable plateau, perfusion was switched to an agonist-containing solution supplemented with the test compound, and changes in current amplitude were continuously recorded. Current inhibition was calculated as inhibition (%) = (1 − I_drug/I_control) × 100, where I_control and I_drug represent the magnitudes of agonist-evoked currents before and during compound application, respectively. Concentration–inhibition curves were fitted by four-parameter nonlinear regression using the “log(inhibitor) vs. response—Variable slope (four parameters)” model in GraphPad Prism to determine half-maximal inhibitory concentrations (IC₅₀).

### Serotonin transporter uptake inhibition assays

The inhibitory activities of compounds against the human serotonin transporter (SERT) were determined using cell-based fluorescent uptake assays. HEK293 cells stably expressing human SERT were seeded into 96-well black clear-bottom plates at 3×10^4^ cells per well and cultured overnight at 37 °C in a humidified atmosphere containing 5% CO₂. On the day of the assay, cells were equilibrated in assay buffer and incubated with serial concentrations of compounds for 30 min at 37 °C. Transporter-mediated uptake was then assessed using a fluorescent uptake detection reagent prepared in HBSS, followed by incubation for 60 min at 37 °C. Fluorescence was measured at an excitation wavelength of 440 nm and an emission wavelength of 520 nm. Centanafadine was used as the positive control, and vehicle-treated wells served as the negative control. Percentage inhibition was normalized to the positive-and vehicle-control responses, and concentration–response curves were fitted by nonlinear regression to determine IC₅₀ values.

### Behavioral experiments

#### Chronic unpredictable mild stress (CUMS)

Mice were subjected to two randomly mild stressors daily for 4 weeks in an unpredictable sequence, including foot shock, ice-water swim, cage tilt, confinement, water or food deprivation, restraint, tail pinch, damp bedding, light-on at night, and novel odorant exposure. No single stressor was applied on consecutive days and stressor sequence varied weekly. Control mice were handled daily and housed under standard conditions without stress exposure. Behavioral experiments were conducted after the onset of CUMS and single administration of drug, followed sequentially by the sucrose preference test and forced swimming test^29^.

#### Chronic restraint stress (CRS) paradigm

Mice were subjected to a chronic restraint stress procedure for 14 consecutive days. Model mice were immobilized in an acrylic restraint tube for 4–6 hours daily during the light phase. Food and water were withheld during the restraint sessions but were available ad libitum at all other times. Control mice were housed under standard conditions without stress exposure. Sucrose preference test was conducted after the onset of CRS and single administration of drug^30^.

### Sucrose preference test (SPT)

Sucrose preference was assessed using a two-bottle choice paradigm^31^. Briefly, mice were first habituated to two bottles containing 1% sucrose solution for 24 h. The following day, one sucrose bottle was replaced with tap water, and mice continued adaptation for an additional 24 h, with the positions of the sucrose and water bottles interchanged at the 12-h time point to control for side bias. Following this 2-day adaptation period, mice were deprived of water overnight. After drug administration, mice were returned to their home cages with two bottles (one containing 1% sucrose solution and the other containing tap water) for a 24-h sucrose preference test. The positions of the two bottles were again swapped at the 12-h mark.Sucrose preference was defined as the ratio of sucrose consumption weight (g) to total liquid consumption.

### Forced swim test (FST)

The FST was performed to evaluate depressive-like behavior in mice. Mice were individually placed into transparent glass cylinders (height: 20 cm, diameter: 15 cm) filled with water (depth: 15 cm) maintained at 22-23℃. The test was recorded by a video camera positioned at the side of the apparatus. Immobility time was quantified from the last 4 min of the 6-min session using Noldus software. Immobility was defined as the absence of active swimming or climbing movements, with the mouse floating passively and making only minimal limb movements necessary to keep its head above the water surface.

### Open field test (OFT)

The open field test was performed to assess spontaneous locomotor activity. Mice were transported to the testing room and allowed to acclimate for at least 30 min before the experiment. Mice were gently placed into the center of an opaque acrylic open field arena (40×40×40 cm) and allowed to explore the apparatus freely for 30 min after administration. Total distance was automatically analyzed by Any-maze software.Between trials, the arena was thoroughly cleaned with 75% ethanol to eliminate olfactory cues that could influence subsequent subjects behavior.

### Conditioned place preference (CPP)

The CPP apparatus was an opaque box divided into three compartments: a drug-paired chamber, a vehicle-paired chamber, and a central transition chamber. The compartments featured distinct floor textures and were separated by removable partitions. Mice behavior was recorded and analyzed using Any-maze software. Entry into a compartment was defined as the animal’s head crossing into the arena.After each trial, all compartments were cleaned with 75% ethanol.

Following acclimation to the housing conditions, mice underwent a 3-day pre-conditioning phase. All partitions were removed, and mice were placed in the apparatus through central transition chamber and allowed to explore freely for 20 min before returned to home cages. This procedure was performed once daily for 3 days. The time spent in each compartment during the 5–20 min interval was recorded, and the preference score was calculated as: (time spent in the compartment / total time spent in both side compartments) × 100%. The compartment with the lower preference score was designated as the drug-paired chamber for the mice.

During the conditioning phase, all the partitions were closed. On days 1, 3, 5, 7, 9, 11, and 13, mice received drug injections (control mice received vehicle injections) and were confined to the drug-paired chamber for 30 min. On days 2, 4, 6, 8, 10, 12, and 14, mice received vehicle injections and were confined to the unpaired chamber for 30 min.

One day after the conditioning phase, the post-conditioning test was conducted. All partitions were removed, and mice were placed in the apparatus through central transition chamber and allowed to explore freely for 20 min. All behavioral tests were performed by experimenters blinded to the treatment groups. The time spent in each compartment was recorded during the 5–20 min interval. CPP score was calculated as the difference of time in drug-paired side between post-test and pre-test^32^.

### Rotarod test

Mice were trained on the rotarod twice daily for 3 consecutive days. The rod rotated at a constant speed of 5 rpm in the first training session. The speed was increased by 5 rpm in each subsequent training session, with each trial lasting 1 min. Mice that fell off the rod before the end of the trial were returned to their home cage and allowed to rest before the next session. Training sessions were spaced at least 2 h apart to allow sufficient recovery between trials. Following rotarod training, mice were screened and randomly assigned into parallel groups based on their screening performance.

Animals failing to remain on the rotarod for at least 110 s during screening were excluded from subsequent analysis. Rotarod tests were conducted at 10, 30, and 90 minutes after drug or vehicle administration. Both the screening and testing conditions involved a linear acceleration from 5 to 40 rpm over a 5-minute period. The latency to fall was recorded as the time when the mouse fell off or rotated with the rod for more than two complete revolutions.

### Statistics

Sample size was referred to previous studies in the same field. In all the experiments, data acquisition and analysis were performed in a double-blinded manner. All the results are presented as the means ±SEM. All statistical analyses were performed using GraphPad Prism software. All data were first tested for normality before statistical testing. Between-group comparisons used unpaired *t* tests or Mann-Whitney test depending on distribution. Welch’s correction was applied to the parametric test in cases of unequal variances between groups. The significance level was set as *P* < 0. 05 (\**P* < 0. 05, \*\**P* < 0. 01, and \*\*\**P* < 0. 001).

## REFERENCES AND NOTES

1 Rincón-Cortés, M. & Grace, A. A. Antidepressant effects of ketamine on depression-related phenotypes and dopamine dysfunction in rodent models of stress. Behavioural Brain Research 379, doi:ARTN 112367 10.1016/j.bbr.2019.112367 (2020).

2 Hirschfeld, R. M. A. Depression Epidemiology and Its Treatment Evolution. J Clin Psychiat 73, E29–E34, doi:DOI 10.4088/JCP.11096tx3c (2012).

3 Global, regional, and national burden of 12 mental disorders in 204 countries and territories, 1990-2019: a systematic analysis for the Global Burden of Disease Study 2019. Fortschr Neurol Psyc 90, 390–391, doi:10.1055/a-1832-0126 (2022).

4 Lucantonio, F., et al. Ketamine Rescues Anhedonia by Cell-Type and Input Specific Adaptations in the Nucleus Accumbens. Biological Psychiatry 97 (2025).

5 Kim, J. W., Suzuki, K., Kavalali, E. T. & Monteggia, L. M. Ketamine: Mechanisms and Relevance to Treatment of Depression. Annual Review of Medicine 75, 129–143, doi:10.1146/annurev-med-051322-120608 (2024).

6 Ma, S. S., et al. Sustained antidepressant effect of ketamine through NMDAR trapping in the LHb (vol 622, pg 802, 2023). Nature 623, E11–E11, doi:10.1038/s41586-023-06814-x (2023).

7 Zanos, P. & Gould, T. D. Mechanisms of ketamine action as an antidepressant. Molecular Psychiatry 23, 801–811, doi:10.1038/mp.2017.255 (2018).

8 Hess, E. M., Riggs, L. M., Michaelides, M. & Gould, T. D. Mechanisms of ketamine and its metabolites as antidepressants. Biochemical Pharmacology 197, doi:ARTN 114892 10.1016/j.bcp.2021.114892 (2022).

9 Cichon, J., et al. Ketamine triggers a switch in excitatory neuronal activity across neocortex. Nature Neuroscience 26, 39-+, doi:10.1038/s41593-022-01203-5 (2023).

10 Jiang, Y. H., Dong, Y. Y. & Hu, H. L. The N-methyl-D-aspartate receptor hypothesis of ketamine’s antidepressant action: evidence and controversies. Philos T R Soc B 379, doi:ARTN 20230225 10.1098/rstb.2023.0225 (2024).

11 Russo, S. J. & Nestler, E. J. The brain reward circuitry in mood disorders. Nature Reviews Neuroscience 14, 609–625, doi:10.1038/nrn3381 (2013).

12 Belujon, P. & Grace, A. A. Restoring Mood Balance in Depression: Ketamine Reverses Deficit in Dopamine-Dependent Synaptic Plasticity. Biological Psychiatry 76, 927–936, doi:10.1016/j.biopsych.2014.04.014 (2014).

13 Cardona-Acosta, A. M. & Bolaños-Guzmán, C. A. Role of the mesolimbic dopamine pathway in the antidepressant effects of ketamine. Neuropharmacology 225, doi:ARTN 109374 10.1016/j.neuropharm.2022.109374 (2023).

14 Fink, K. B. & Göthert, M. 5-HT receptor regulation of neurotransmitter release. Pharmacol Rev 59, 360–417, doi:10.1124/pr.107.07103 (2007).

15 Peters, K. Z., Cheer, J. F. & Tonini, R. Modulating the Neuromodulators: Dopamine, Serotonin, and the Endocannabinoid System. Trends in Neurosciences 44, 464–477, doi:10.1016/j.tins.2021.02.001 (2021).

16 Molinari, M., et al. Dopamine and serotonin cotransmission filters striatonigral synaptic activity via 5-HT1B receptor activation. Sci Adv 11, doi:ARTN eadx4577 10.1126/sciadv.adx4577 (2025).

17 Simmler, L. D., et al. Dual action of ketamine confines addiction liability. Nature 608, 368-+, doi:10.1038/s41586-022-04993-7 (2022).

18 Alex, K. D. & Pehek, E. A. Pharmacologic mechanisms of serotonergic regulation of dopamine neurotransmission. Pharmacology & Therapeutics 113, 296–320, doi:10.1016/j.pharmthera.2006.08.004 (2007).

19 Zhou, J. S., et al. Activation of postsynaptic 5-HT1A receptors improve stress adaptation. Psychopharmacology 231, 2067–2075, doi:10.1007/s00213-013-3350-z (2014).

20 Nakatsuka, D., et al. Fine-tuning of dopamine receptor signaling with aripiprazole counteracts ketamine’s dissociative action, but not its antidepressant effect. Translational Psychiatry 15, doi:ARTN 77 10.1038/s41398-025-03284-9 (2025).

21 Wan, L. B., et al. Ketamine Safety and Tolerability in Clinical Trials for Treatment-Resistant Depression. J Clin Psychiat 76, 247-+, doi:10.4088/JCP.13m08852 (2015).

22 Smith-Apeldoorn, S. Y., Veraart, J. K. E., Spijker, J., Kamphuis, J. & Schoevers, R. A. Maintenance ketamine treatment for depression: a systematic review of efficacy, safety, and tolerability. Lancet Psychiat 9, 907–921 (2022).

23 Acero-Castillo, M. C., et al. Is the antidepressant effect of ketamine separate from its psychotomimetic effect? A review of rodent models. Neuropharmacology 258, doi:ARTN 110088 10.1016/j.neuropharm.2024.110088 (2024).

24 Kokkinou, M., Ashok, A. H. & Howes, O. D. The effects of ketamine on dopaminergic function: meta-analysis and review of the implications for neuropsychiatric disorders. Molecular Psychiatry 23, 59–69, doi:10.1038/mp.2017.190 (2018).

25 Sawyer, E. K., et al. Neurobiological Changes Mediating the Effects of Chronic Fluoxetine on Cocaine Use. Neuropsychopharmacology 37, 1816–1824, doi:10.1038/npp.2012.29 (2012).

26 Ago, Y., et al. (R)-Ketamine Induces a Greater Increase in Prefrontal 5-HT Release Than (S)-Ketamine and Ketamine Metabolites via an AMPA Receptor-Independent Mechanism. Int J Neuropsychoph 22, 665–674, doi:10.1093/ijnp/pyz041 (2019).

27 Ago, Y., Yokoyama, R., Asano, S. & Hashimoto, H. Roles of the monoaminergic system in the antidepressant effects of ketamine and its metabolites. Neuropharmacology 223, doi:ARTN 109313 10.1016/j.neuropharm.2022.109313 (2023).

28 Zhuo, Y. Z., et al. Improved green and red GRAB sensors for monitoring dopaminergic activity in vivo. Nature Methods 21, doi:10.1038/s41592-023-02100-w (2024).

29 Zhang, Y. F., et al. Ventral striatal islands of Calleja neurons bidirectionally mediate depression-like behaviors in mice. Nature Communications 14, doi:ARTN 6887 10.1038/s41467-023-42662-z (2023).

30 Yue, C. Y., et al. Adenosine signalling drives antidepressant actions of ketamine and ECT. Nature 649, doi:ARTN 410462 10.1038/s41586-025-09755-9 (2026).

31 Willner, P., Towell, A., Sampson, D., Sophokleous, S. & Muscat, R. Reduction of Sucrose Preference by Chronic Unpredictable Mild Stress, and Its Restoration by a Tricyclic Antidepressant. Psychopharmacology 93, 358–364 (1987).

32 Yousefzadeh, S. A., et al. A cholinergic hub in the nucleus accumbens gates opioid-reward learning. Nature 658, doi:10.1038/s41586-026-10887-9 (2026).

